# Help thine enemy: the evolution of short ranging signals

**DOI:** 10.1101/255836

**Authors:** Szabolcs Számadó

## Abstract

The proximity risk model offers one possible explanation of honest signalling of aggressive intent in biology. This model assumes that the probability of successful attack is a function of the distance between the contestants and that this distance can be correctly estimated. This later assumption may not hold in nature where contestants have to estimate this distance under noisy conditions. Here I investigate with the help of a game theoretical model whether short-range ranging signals can be evolutionarily stable under such conditions. These signals can help the opponent to estimate the correct distance, thus they can promote honest signalling of intentions. Here I show that ranging signals that help the estimation of distance between opponents can be evolutionarily stable. However, such help only benefits those individuals who are able and willing to attack. As a result, ranging signals in themselves are an honest cue of proximity and in turn they are honest cues of aggressive intent. I give an example: “soft-song” in birds, and I discuss the predictions of the model.

## 1. Background

The evolution of aggressive communication is challenging to explain. The difficulty stems from the fact that there is a conflict of interest between participants as aggressive communication is usually tied to resource competition [1, 2]. Since conflict of interest promotes dishonest signalling the evolution of honest signals seems to be unlikely under such scenario [1, 2]. Why would individuals reveal their state when they could gain advantage by misinformation? Enquist [3] was the first to show in his seminal model that communication of states or intentions (depending on the interpretation of the model) can be evolutionarily stable in the context of aggressive communication. Contrary to earlier suggestions that honesty would require some short of “retaliation”, Enquist [3] was able to show that there is no need for it. The condition of honesty is very simple: the potential risk from using dishonest signals have to be larger than the potential benefit resulting from the use of same signals. One crucial condition in the model is that potential cheaters should not be able to flee the conflict even if they want to do so. The reason behind this assumption was unexplained in the original formulation. Later ‘proximity risk’ was proposed as a potential proximate mechanism by Számadó [4]. The idea of the proximity risk model is that there is a distance within which potential cheaters cannot get away with bluffing even if they want to do so, and this is the distance within which threat displays are expected to be honest [4].

Számadó’s [4] model assumes that contestants are able to judge the distance perfectly. However, this is rarely the case in nature: errors of estimation are expected to happen. Such errors in turn might decrease the fitness of an individual if it picks the wrong strategy as a function of the distance. This applies to the opponent as well, in fact errors of the opponent might decrease the fitness of the ego. Here I propose that it could be the interest of the ego to help the opponent to estimate the distance correctly, in order to avoid costly mistakes of the opponent. I call such signals as ‘short-range ranging signals’ (SRRS) since the function of these signals is to help with the estimation of range between opponents. Please note, that the form and function of these signals can be different than the “traditional” ranging signals investigated in the context of territorial bird song (i.e. long-distance broadcast displays [5-8]). I investigate why and under what conditions would such help benefit the signaller and thus under what conditions are short-range ranging signals evolutionarily stable.

## 2. Methods

Enquist’s [3] model is a game of aggressive communication where two individual competes for an indivisible resource. The game has three stages: (i) at the first stage Nature picks the state of the contestants, which can be weak (W) or strong (S); (ii) at the second stage the contestants can chose between two cost-free signals A or B; finally (iii) at stage three the contestants can either flee (F), attack (A) or wait for the opponent to flee and attack only if the opponent stays to fight (cond.A). Figure 1 depicts the structure of the game (after [9]). Let *V* denote the value of the resource. The cost of fighting depends on the strength of the ego and that of the opponent; accordingly, there are four cost parameters: *C*_*WW*_, *C*_*WS*_, *C*_*SW*_ and *C*_*SS*_. The following relation is assumed to hold between these parameters: *C*_*WS*_ > *C*_*WW*_, *C*_*SS*_ > *C*_*SW*_. On top of the cost of fighting there are other potential costs as well: (i) attacking a fleeing opponent (*F*_*A*_), (ii) pausing when the opponent attacks (*F*_*p*_), and finally there is a cost to flee (*F*_*f*_). Table 1 sums up the pay-offs of the game.

**Table 1.**
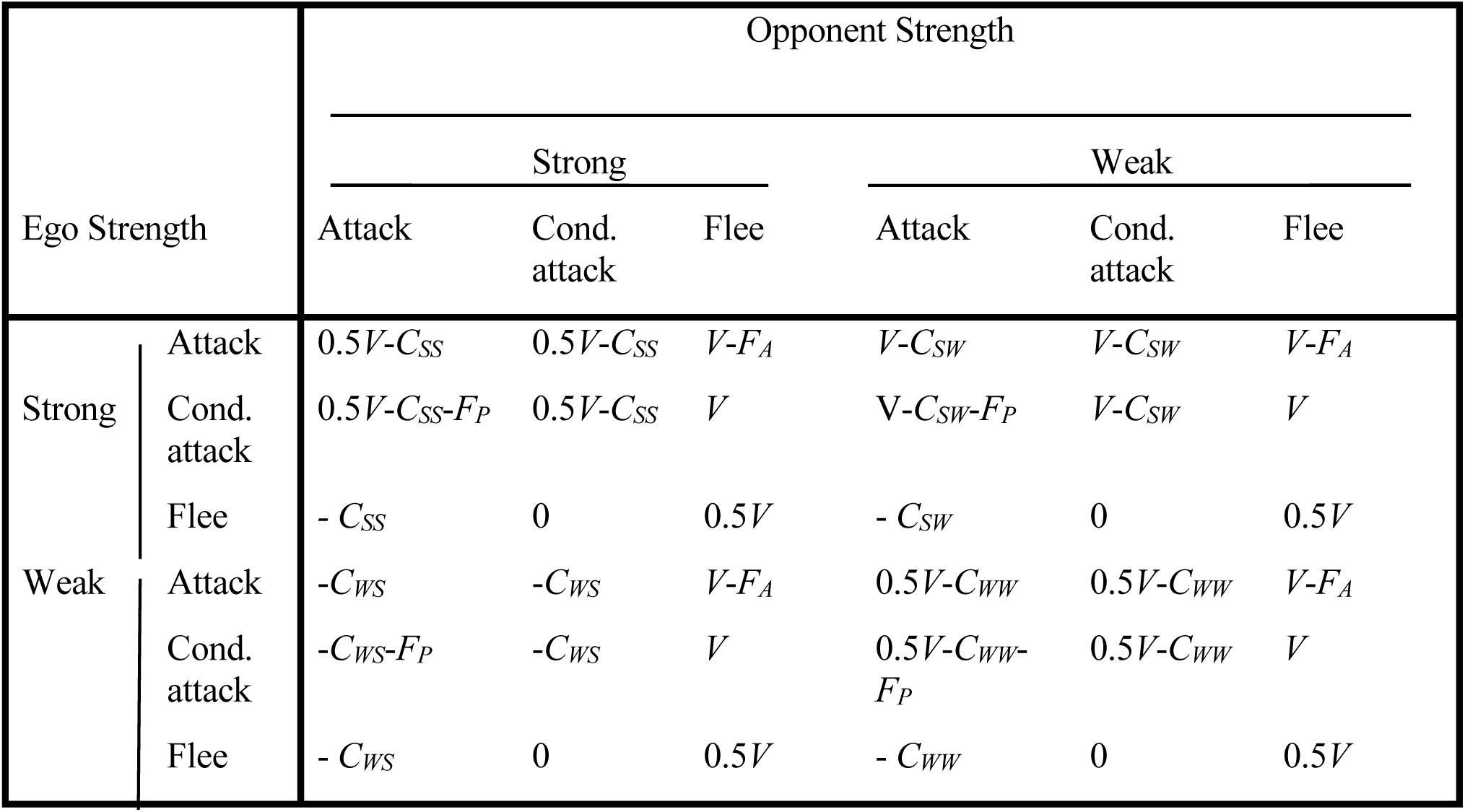
Payoffs matrix (modified after [10]). *V*: amount of the contested resource; *C*_*SS*_, *C*_*WW*_: expected cost of fight between equal opponents; *C*_*SW*_: cost payed by strong individual to beat weak one; *C*_*WS*_: cost payed by weak individual when beaten by strong one; *F*_*f*_: cost of fleeing; *F*_*A*_: cost of attacking fleeing opponent; *F*_*P*_: cost of waiting if the opponent attacks unconditionally.

| Ego Strength |  | Opponent Strength |  |  |  |  |  |
| --- | --- | --- | --- | --- | --- | --- | --- |
|  |  | Strong |  |  | Weak |  |  |
|  |  | Attack | Cond.<br>attack | Flee | Attack | Cond.<br>attack | Flee |
| Strong | Attack | $0.5V - C_{SS}$ | $0.5V - C_{SS}$ | $V - F_A$ | $V - C_{SW}$ | $V - C_{SW}$ | $V - F_A$ |
| | Cond.<br>attack | $0.5V - C_{SS} - F_P$ | $0.5V - C_{SS}$ | $V$ | $V - C_{SW} - F_P$ | $V - C_{SW}$ | $V$ |
| | Flee | $-C_{SS}$ | 0 | $0.5V$ | $-C_{SW}$ | 0 | $0.5V$ |
| Weak | Attack | $-C_{WS}$ | $-C_{WS}$ | $V - F_A$ | $0.5V - C_{WW}$ | $0.5V - C_{WW}$ | $V - F_A$ |
| | Cond.<br>attack | $-C_{WS} - F_P$ | $-C_{WS}$ | $V$ | $0.5V - C_{WW} - F_P$ | $0.5V - C_{WW}$ | $V$ |
| | Flee | $-C_{WS}$ | 0 | $0.5V$ | $-C_{WW}$ | 0 | $0.5V$ |

**Figure 1.**
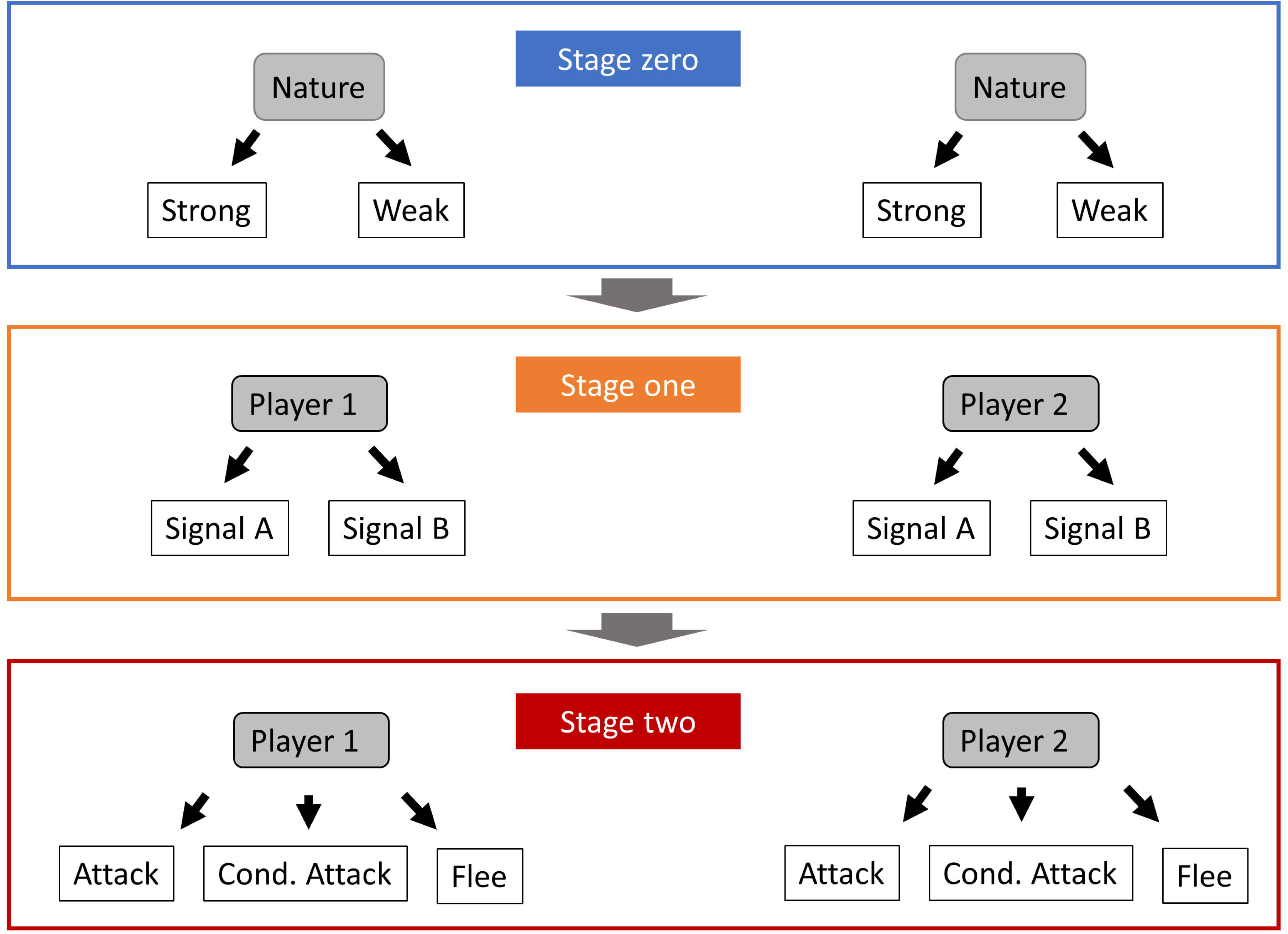
Schematic description of Enquist’s game [3] (modified after Számadó [9]). Stage zero: Nature picks the state of the contestants; this stage is hidden from other players. Stage one: each contestant picks a signal, A or B. Stage two: each contestant picks a behaviour as a response to the signal: Flee, Conditional Attack or Attack.

While in Enquist’s [3] game an attack was always successful, Számadó [4] investigated a scenario where the chance of a successful attack depends on the distance between the participants. Let *x* denote the distance between the opponents and *f*(*x*) denote the probability of successful attack a function of distance. The function *f*(*x*) is assumed to be monotonically decreasing function of *x*, i.e. the further is the opponent the smaller is the chance to strike it successfully. Számadó [4] assumed that individuals know the distance separating the opponents (*x\**). Here I assume that they estimate this distance (*x*_*e*_) and this estimation has an error *φ* (0, *σ*) normally distributed, where *σ* denotes the standard deviation.

Enquist [3] defined the honest global strategy denoted *S*_*A*_ as follows (p. 1155):

„ *If strong, show A; if the opponent also shows A attack and if the opponent shows B, repeat A and attack only if it does not withdraw immediately. If weak show B and give up if the opponent shows A and attack if the opponent shows B*.”

Enquist [3] investigated the evolutionary stability of this strategy *S*_*A*_ against a cheater type in which weak individuals show A instead of B. The corresponding global strategy (*S*_*B*_) can be written up as follows [10] (p. 222):

„*Display always A in the first round, regardless of strength; then in the second round if strong attack unconditionally if opponent shows A or wait until opponent flees if it has shown B; if weak withdraw if opponent signals A or wait until opponent flees if it has shown B.*”

Here we assume that the strategy of an individual is defined as playing the honest strategy (*S*_*A*_) with probability 1-*p*(*x*) and playing *S*_*B*_ with probability *p*(*x*) as a function distance (*x*).

## 3. Results

Here, I investigate the evolutionary stability of ranging signals using the proximity risk model proposed by Számadó [4]. First I recapture the key results of previous models. Then I calculate the ESS condition for short-range ranging signals. Finally I give and interpretation and an example of the result.

### Proportion of cheaters and distance thresholds in the proximity risk model

There is a region in the proximity risk model where a mixed strategy (*S*_*M*_) is an evolutionarily stable strategy (ESS). This mixed ESS is supported by cheater (*S*_*B*_) and honest (*S*_*A*_) pure strategies, where the probability of playing these strategies are *p* and 1-*p* respectively. The equilibrium ratio of mixing is given by the following equation [4]:

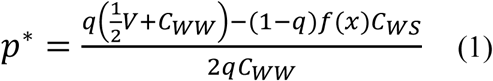

Starting from this equation one can calculate the threshold distance between honest signalling and mixed cheating (*x*_*HSD*_) where *p\**=0, as well as the threshold between mixed cheating and no signalling (*x*_*DSD*_, see [4]) where *p*\*=1:

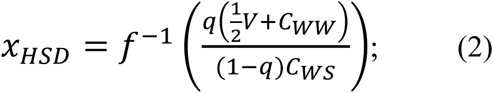

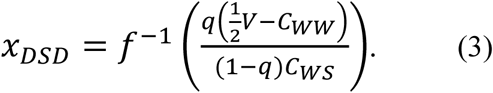

Figure 2 depicts an example. The blue line shows the threshold below which honesty is the ESS whereas the red line shows the threshold above which there is no honest communication. The mixed strategy *S*_*M*_ is an ESS between these thresholds. Figure 3 shows the equilibrium proportion of cheaters (*p\**) in this region (see [4]).

**Figure 2.**
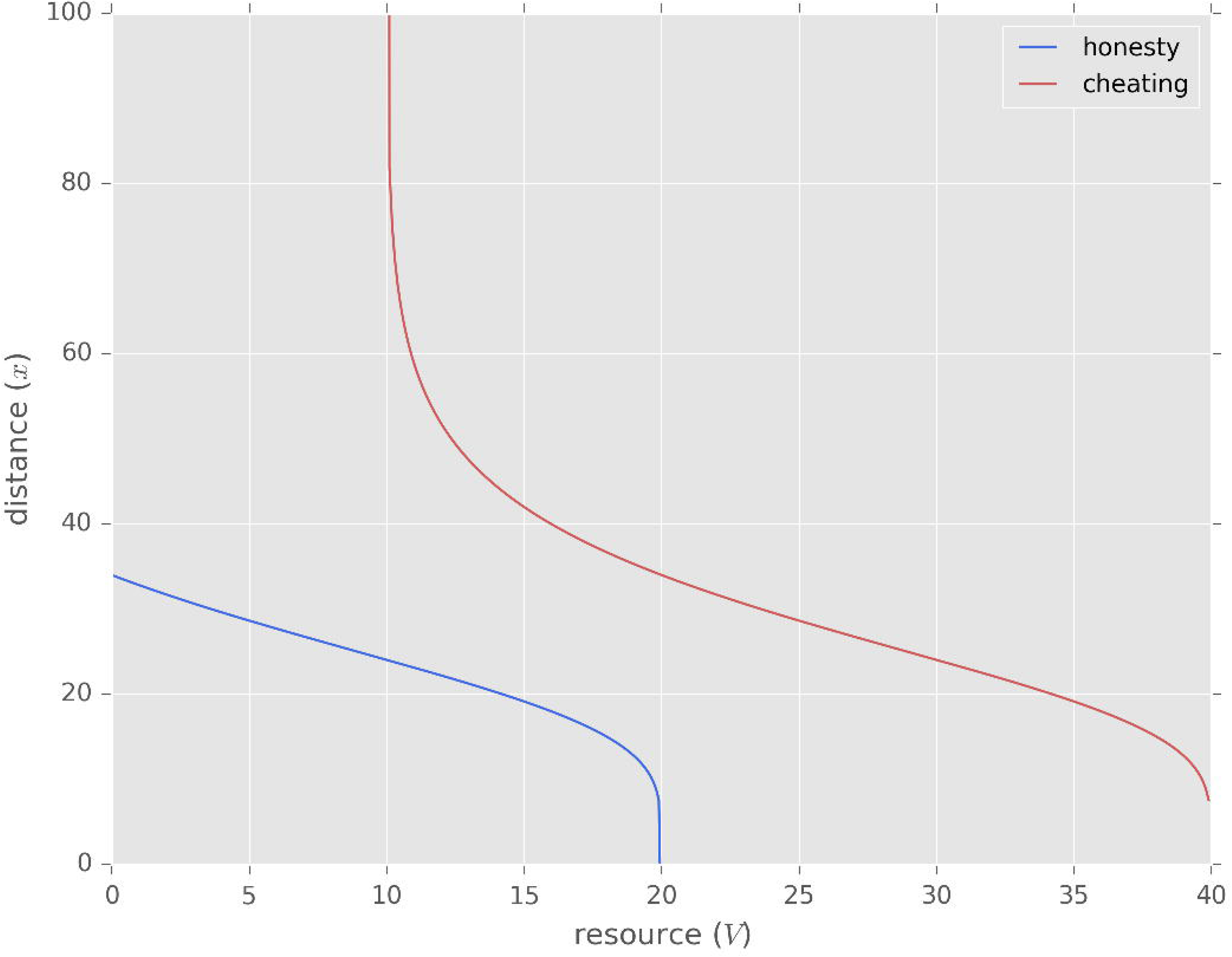
Honest, mixed cheating and dishonest regions in spatially explicit game of aggressive communication. Blue line (*x*_*HSD*_): threshold between honest (*S*_*A*_) and mixed cheating (*S*_*M*_); red line (*x*_*DSD*_): threshold between mixed cheating (*S*_*M*_) and dishonest region (only *S*_*B*_).

### ESS conditions of ranging signals in the proximity risk model

Let’s assume that estimating the distance without any help is prone to errors. Let’s further assume a population where individuals give a ranging signal (*Rs*) to help the opponent to estimate distance *x*. At the equilibrium both opponent will use this signal; both of them will estimate the distance correctly, thus they both will play *p\**. Can this equilibrium be invaded by a mutant strategy that does not use *Rs*? The use of *Rs* to be ESS the following condition must hold [11]:

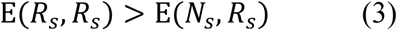

if E(R_s_, R_s_) = E(N_s_, R_s_) then

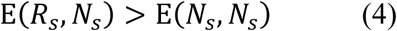

Since opponents of individuals using *Rs* can estimate distance correctly (thus they will play *p\**), whereas opponents of individuals not giving *Rs* cannot do so (hence will play *p* ≠ *p*\*), Eq.3 implies:

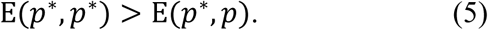

Then the equilibrium condition (Eq. 5) can be expressed with these terms as follows:

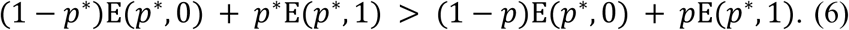

After rearrangement the ESS condition of short-range ranging signals is as follows:

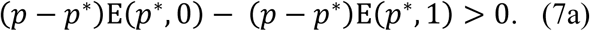

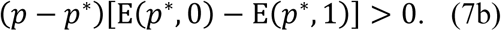

Where the fitness of an individual playing *p*\* against individuals playing the honest strategy (*S*_*A*_) is as follows:

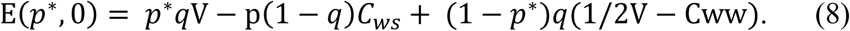

The fitness of an individual playing p* against individuals playing the cheater strategy (*S*_*B*_) is as follows:

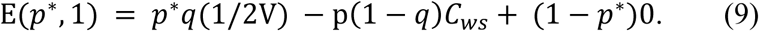

### Interpretation and examples

There are two obvious cases: (i) either *p*>*p\**, then the following inequality has to hold:

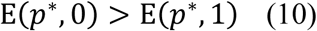

That is, if the proportion of cheaters is higher as specified by the mixed strategy (*p*>*p\**), then individuals playing the mixed strategy against the honest type has to do better than individuals playing the mixed strategy against the cheater type. This condition will hold as long as:

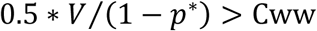

(ii) In the second case *p*<*p\**, then the following inequality has to hold:

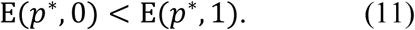

That is, if the proportion of cheaters is lower as specified by the mixed strategy (*p*<*p\**), then individuals playing the mixed strategy against the cheater type has to do better than individuals playing the mixed strategy against the honest type.

Let’s define ΔE(p^*^) as a difference in fitness when playing against an opponent playing the equilibrium probability of mixing and an opponent playing an out-of-equilibrium mixing:

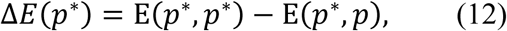

Which in turn is the left-hand side of Eq. 7, thus:

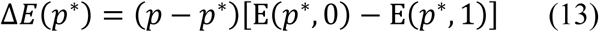

Let’s define ΔE as a fitness difference between playing honest and dishonest opponent, while the ego plays *p\**:

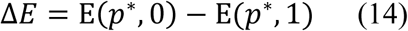

Finally, let’s further define Δp as follows:

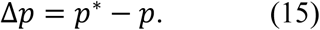

ΔE can be calculated by substituting Eqs. 8, 9 and it results the following equation:

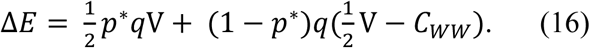

Figure 4 shows an example for the error of estimating the difference between opponents (*x*_*e*_). Figure 5. A shows an example for ΔE. It is clear that ΔE is always positive, which means that it is always more beneficial to play against the honest type than the cheater type. Fig 5.B shows the change in the frequency of cheaters (Δp) as a result of this error of estimation. It is clear that this error of estimation has the greatest effect in the region where the mixed strategy is the ESS. Fig 5.B also shows that the proportion of cheaters is increased at close range and decreased at mid-range compared to the ESS mixing ratio. Combining the insights of Fig. 5.A and 5.B predicts that if opponents underestimate the probability of a successful attack (due to overestimating distance, e.g. Fig. 5.B close distance), thus they are likely to cheat with a higher probability than the ESS mixing ratio, then it is the interest of the ego to help them to make a correct estimation so that the opponent’s choice to play the honest type increased, which is beneficial to the ego (i.e. Fig.5.B). Figure 5. C, shows this effect, i.e. it shows ΔE(p^*^). There are two regions where playing *p\** has higher fitness than a mutant (i.e. regions where *p\** is an ESS): (i) the first region is at low value of resource and large distance; (ii) whereas the second region is at high value of resource and low distance. Assuming that territorial disputes represent high value of resource and individuals are aware of the fact that they are fighting for a territory, it is clear that the use of ranging displays (*Rs*) is an honest cue of proximity in this region. In other words, individuals receiving a ranging display from the opponent contesting a resource of high value can be certain that the opponent is close enough to strike with a high success rate.

**Figure 4.**
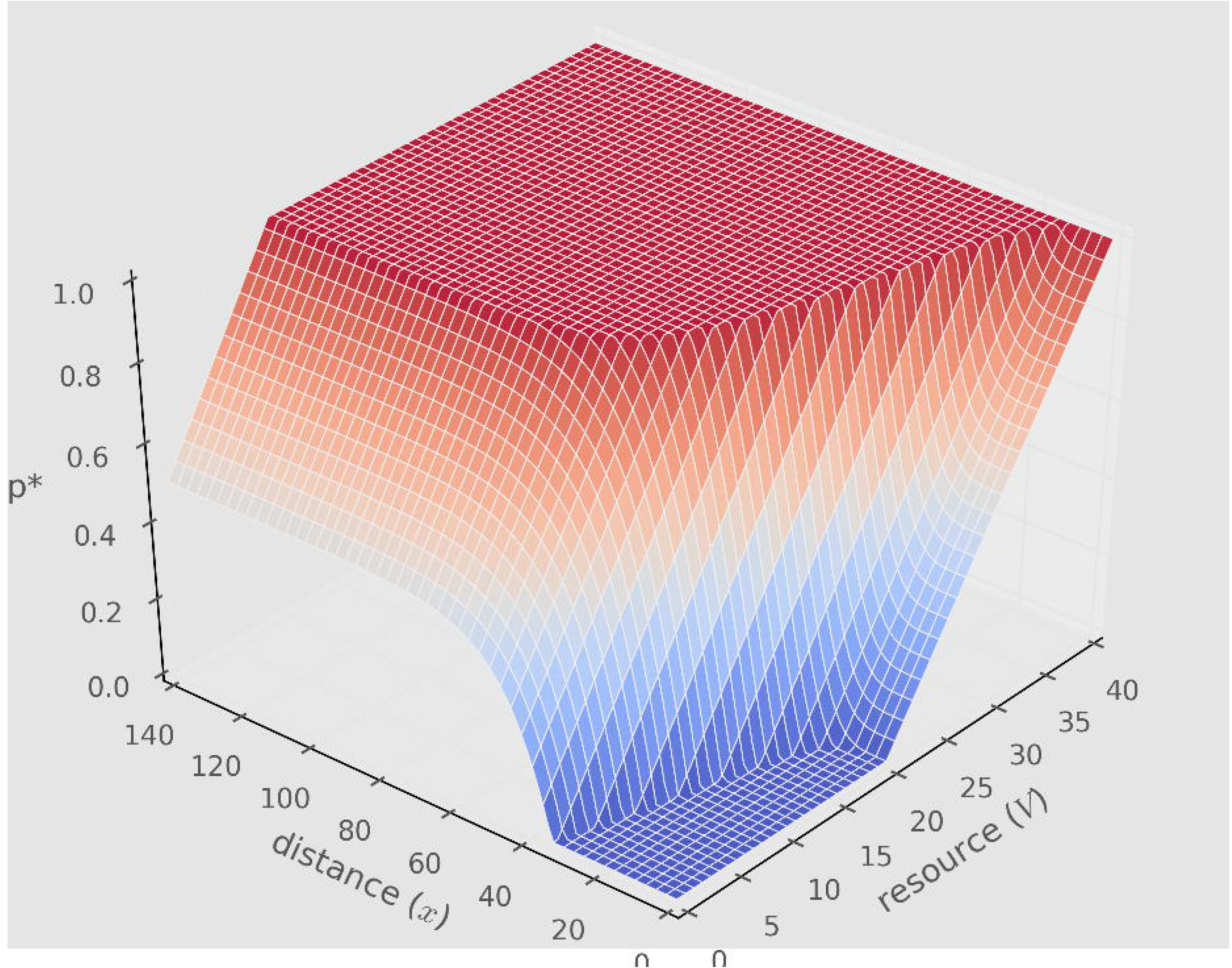
The probability of successful attack (*f*(*x*)) as a function of distance (*x*) without and with error of estimation (*σ*), where *f*(*x*) function is described by a Gompertz curve: 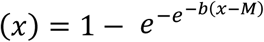; where *b* = 0.1, *M* = 25.

**Figure 5.**
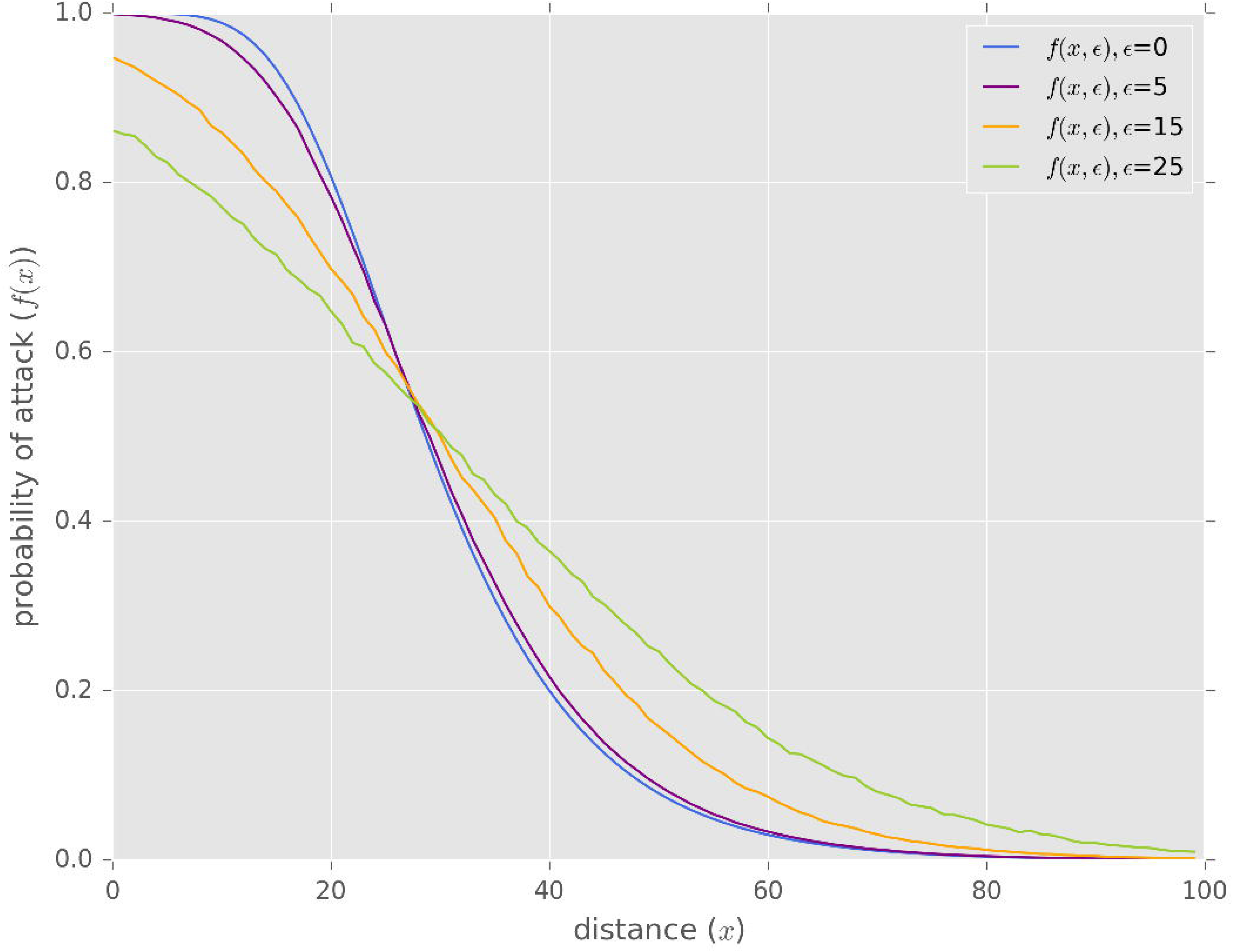
(a) The difference in fitness when playing against an honest opponent vs. playing against a cheater (ΔE). (b) The change in the probability of cheating (Δp) as a result of the error of estimating the difference as a function of distance (*x*) and the value of the resource (*V*). (c) The difference in fitness (ΔE(p^*^)) when using a ranging signal (*Rs*) vs. not using (*Ns*) in a population of individuals using *Rs*.

Figure 6 shows examples of Δp (Fig.6. A,B and C) and the corresponding ΔE(p^*^) (Fig.6. D,E and F) values for different estimates of error (A,D *σ* =5; B,E *σ* =15, C,F *σ* =25). It clear that error prone estimation of distance mostly affects the zone of mixed cheating (i.e. the zone between the dashed lines). The larger the error the larger this zone. Also it starts to have an effect on the honest zone at higher resource values (Fig 6.F). So far it was assumed that ranging signals are cost-free. However, introducing a small cost (*c*_*R*_) into Eq.7 (left-hand side) makes the first region (low value of resource, high distance) mostly disappear (depending on the value of cost and on the value of error). Figure 6.G, H and I show an examples. Thus, ranging signals are honest cues of proximity regardless of the value of resource under such scenario (e.g. Fig 6.H). Figure 6. G also shows that the use of *Rs* might not be an ESS when *Rs* is costly if the error of estimating the distance is small.

**Figure 6.**
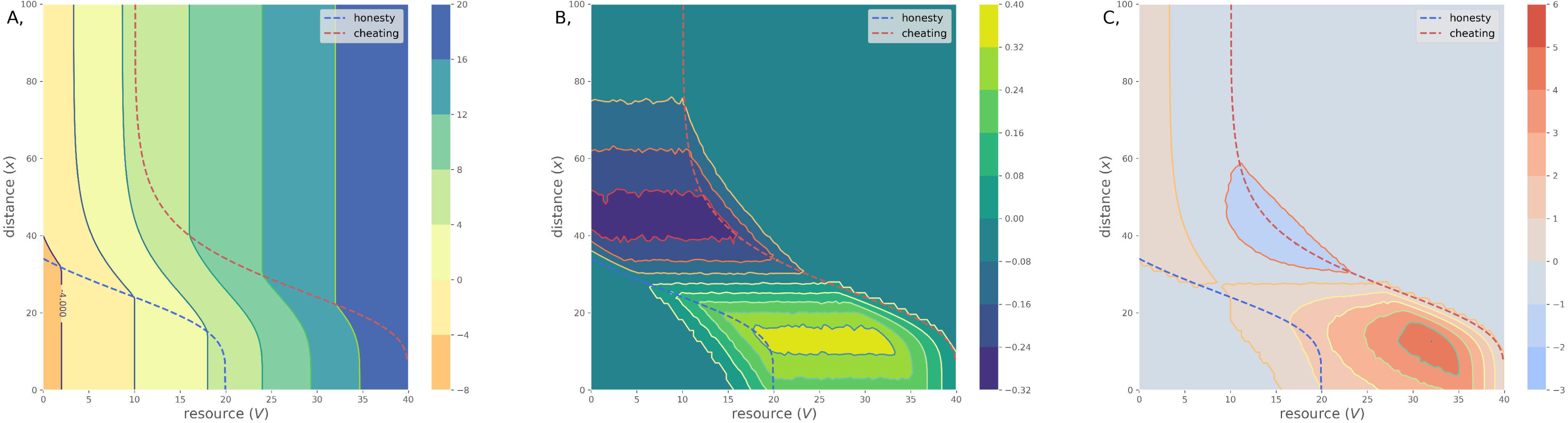
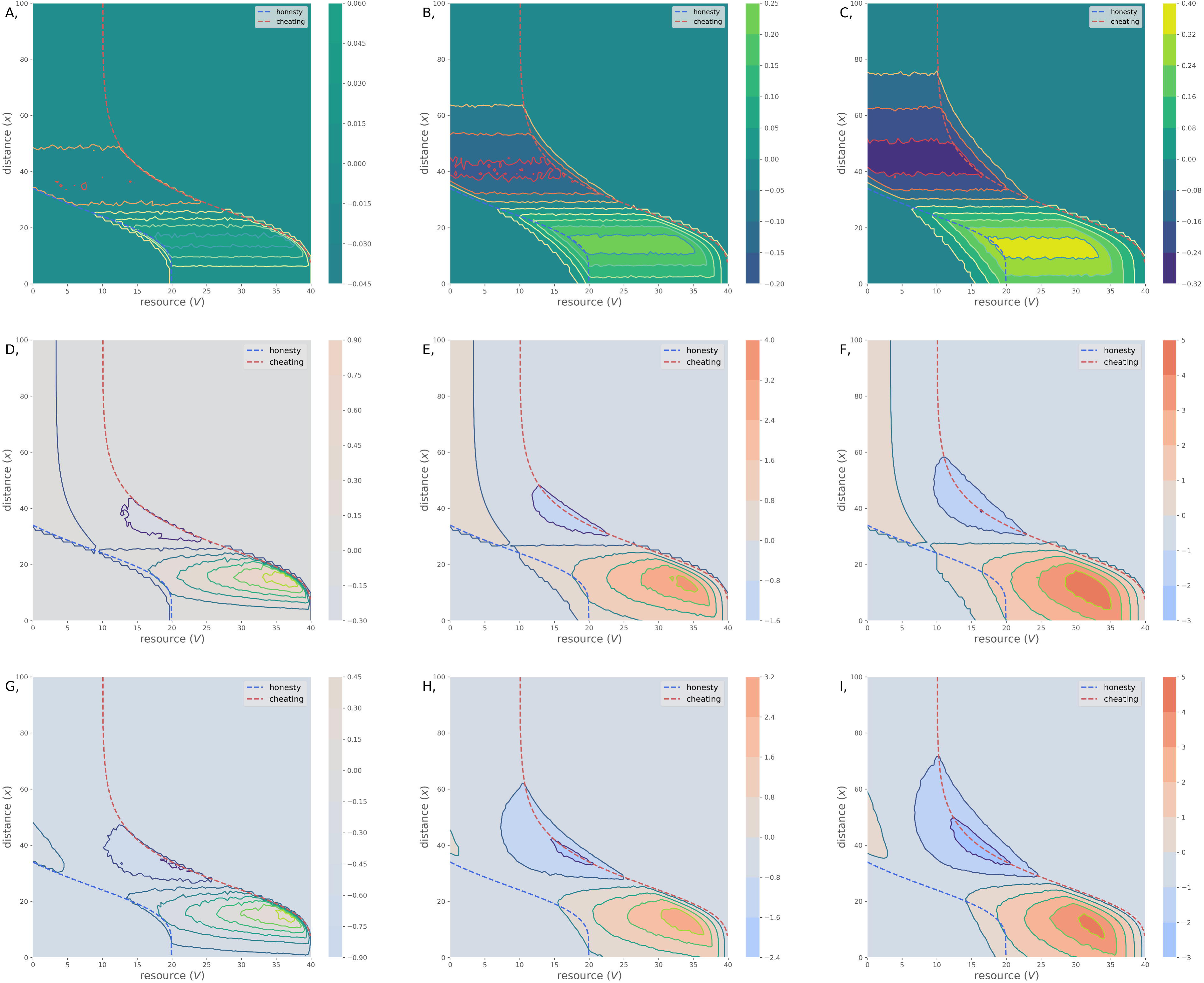
First row: the change in the probability of cheating as a result of the error of estimating the difference as a function of distance (*x*) and the value of the resource (*V*) as a function of different errors of estimation: (A) *σ*=5, (B) *σ*=15, and (C) *σ*=25. Second row: the difference in fitness (ΔE(p^*^)) when using a ranging signal (*Rs*) vs. not using (*Ns*) in a population of individuals using *Rs* as a function of different errors of estimation: (D) *σ*=5, (E) *σ*=15, and (F) *σ*=25. Third row: the difference in fitness (ΔE(p^*^)) when using a ranging signal (*Rs*) vs. not using (*Ns*) in a population of individuals using a costly signal where *c*_*R*_= 0.5; different errors of estimation: (G) *σ*=5, (H) *σ*=15, and (I) *σ*=25.

## 4. Discussion

I have shown that the use of ranging signals is evolutionarily stable when there is an error of estimating the distance between the two opponents. Cost-free ranging signals are an honest cue of aggressive intentions when the (i) value of the resource is low and the distance between the opponents is high, or the (ii) value is high and the distance is low (Fig. 5.C). Assuming that animals are aware of the value of resource they are competing for then the use of ranging signal is an honest cue of low distance in case of high value of contested resource. A small production cost of ranging signals eliminates the first region; thus, the use of ranging signal is an honest cue of low distance regardless of the value of the resource (Fig. 6.H). The intuitive interpretation of this result is that uncertainty of estimating the distance benefits cheaters, thus only honest individuals are expected to reduce it. Accordingly, if I see my opponent *to help me* to reduce the uncertainty regarding the estimation of distance between the two of us then it is honest cue of my opponent’s aggressive intentions.

Please note that the model’s assumption and thus the current logic applies to all situations where individuals have some error prone estimation of the distance. This estimation need not be vocal only, so that the opponents can have a visual line of sight, yet it would not exclude the current mechanism as long as the visual judgement of distance is not perfect.

“Soft songs” are low-amplitude songs or calls observed in a number of species, mostly in aggressive context [12]. Empirical studies shown that out of a number of song types associated with aggressive context, i.e. song type switching, song matching, soft-song [13] only soft song predicts the probability of attack. Soft song is a reliable predictor of attack in many species including song sparrows (*Melospiza melodia*) [14-17], in swamp sparrows (*Melospiza georgiana*) [18], in brownish-flanked bush warbler (*Cettia fortipes*) [19] and in black-throated blue warblers (*Dendroica caerulescens*) [20]. Soft call, a low amplitude call, which appears to be the equivalent of soft song is a reliable signal of aggression in corncrakes (*Crex crex*) [21, 22]. There has been a long debate on the functions of soft song and whether soft song is the best suited for short-range ranging or not [13, 18, 23, 24]. There is no room to give justice to this debate here. Accordingly, the goal is not to decide the debate, just to point out that from a purely theoretical point of view soft-song fits the function of SRRS proposed in the model. Whether this is the case in nature or not, is an empirical issue.

First, I review the predictions of the model, then I evaluate these predictions in the light of the empirical evidence on soft-song. The current model makes five predictions: (1) SRRS functions in the context of aggressive communication. (2) SRRS is a reliable predictor of attack. (3) SRRS functions in a noisy environment. (4) SRSS functions to reduce the level of noise. (5) The use of SRSS is evolutionarily stable (i) when the value of the resource is low and the distance between the opponents is high, or (ii) when the value of the resource is high and the distance is low; additionally a small cost of displays can eliminate the first region.

(1) SRRS functions in the context of aggressive communication. The majority of empirical examples of soft song is in the context of aggressive communication ([13-21]). This prediction is strongly supported.

(2) SRRS is a reliable predictor of attack. This prediction has a strong support as well. Soft song is the most reliable indicator of attack ([13, 14, 18]).

(3) SRRS functions in a noisy environment. There is no direct evidence pro or contra for this prediction. In general both empiricist [25, 26] and theoreticians [27] agree that communication in nature expected to be noisy. Close distance or line of sight between opponents need not imply that there is no noise. Most bird species have relatively narrow binocular vision. The primary function binocular vision in birds is debated [28] and some argue that that not all bird species might have stereopsis (depth perception) even if they have binocular vision [28, 29]. While Emberizid sparrows (*Emberizidae*) including song sparrows (*Melospiza melodia*) has a relatively wide binocular field [30] compared to other birds, the authors [30] suggest that the main function of binocular vision is contrast recognition, guiding of bill movement and precision grip. If so, simply being in close range and having a line of sight need not resolve the uncertainty of the estimation of distance between opponents.

(4) SRSS functions to reduce the level of noise, i.e. it serves as a signal of proximity. This is the most debated point (see [18, 23] vs. [24]). Yet, this seems to be an unresolved debate, because there is no strong empirical evidence offered pro or contra. There is only a short verbal argument presented in Ballentine at al. [18] which questions whether soft-song is the best choice at close range to reduce uncertainty (p. 700): “Because of the way that sound attenuates with distance, a song that is of low amplitude when it reaches the receiver may be a soft song produced nearby or a broadcast song produced at a greater distance, whereas a song that is of high amplitude when it reaches the receiver is unambiguously a song produced in close proximity.” Unfortunately, this claim is being repeated, and presented as evidence, in subsequent publications without any empirical evidence (Searcy et al. [23]; p. 1215): “Loud song produced close to a receiver, however, is actually a less ambiguous signal of proximity than is a low-amplitude song (Ballentine et al. 2008). Because all sounds attenuate with distance, a song that is still loud when it reaches the receiver must have been produced nearby, whereas a song that is of low amplitude when it reaches the receiver could be a soft song produced nearby or a louder one produced further away.” On the other hand, a recent empirical result found that the structure of soft song is adopted to short range communication [31]; the authors concluded that “both the acoustic structure and low amplitude of this song mode are adapted to reduce transmission range” [31] (p. 119). While the authors interpreted this result in term of the eavesdropping, this result is equally consistent with, and is an expected outcome of the recent proposal.

(5) The stability conditions of SRRS I.: signal cost. First of all, there is no empirical evidence on the fitness cost of using soft-song. The general *assumption* is that the production cost is low [15, 18, 32]. The whole soft-song “paradox” started out because soft-song seems to be a cheap yet reliable signal, which runs contrary to predictions of the Handicap Principle or “costly signalling theory” [33, 34], which is still the dominant paradigm of honest signalling in biology. As Anderson and her colleagues ([32], pp. 1273) write: „It is not clear what if any cost acts to maintain the reliability of soft song as a signal of aggressive intent in song sparrows”. Akcay at al. [17] ask the same question: “So how can a signal that is apparently not costly to produce be a reliable aggressive signal?” (pp. 380). “Receiver-retaliation rule” or “Receiver-dependent cost” was proposed as a solution to this problem [15, 17, 18, 24], but no details and no model was offered as how such “retaliation-cost” would work. The current model can highlight these missing details and it can explain how a low production cost signal of aggressive intent can be reliable and evolutionarily stable.

(5) The stability conditions of SRRS II.: value of the resource and distance between opponents. These points seems to be much less controversial than the previous ones. Most examples of soft-song observed in territorial disputes [13-21] –i.e. high value of resource- and at close range. These conditions fit the second region identified by the current model.

All in all, helping the opponent to make a correct judgment of distance can be beneficial for the ego. Giving such signal is a sign of confidence and aggressive intentions since the reduction of uncertainty only benefits an honest signaller. Signals that evolved for this function expected to have reduced range.

## List of abbreviations

ESS: evolutionarily stable strategy
HSD: honest striking distance
DSD: dishonest striking distance.

## Ethics approval and Consent to Participate

Not applicable.

## Consent for publication

Not applicable.

## Availability of Data and Materials

Not applicable.

## Founding

S.S. was supported by the National Research, Development and Innovation Office (NKFIH) OTKA grant K 108974, by the European Research Council (ERC) under the European Union’s Horizon 2020 research and innovation programme (grant agreement number 648693) and by GINOP 2.3.2-15-2016-00057 (Az evolúció fényben: elvek és megoldások).

## Acknowledgements

Not applicable.

## Author contribution

S.S. conceived the idea, analysed the model and wrote the paper.

